# Structural studies of mycobacterial HptG reveal a silent state and offer insights into TLR4 activation

**DOI:** 10.64898/2025.12.11.693645

**Authors:** Giovanni Barra, Marina Sala, Maria Carmina Scala, Pietro Campiglia, Hwa-Jung Kim, Alessia Ruggiero, Rita Berisio

## Abstract

HtpG of *Mycobacterium tuberculosis* (HtpG_Mtb_) is an ATP-dependent chaperone that assists the correct folding of nascent and stress-accumulated misfolded proteins, in concert with other chaperones. Beside playing a role in stress response, it is able to elicit an immune response against *M. tuberculosis* infection by activating Dendritic Cells in a TLR4-mediated manner. However, we lack a full understanding of the molecular determinants of HtpG_Mtb_ catalytic activity and TLR4 activation, due to the lack of structural and biophysical data. Here, we report the first crystal structure of HtpG_Mtb_, in complex with the non-hydrolysable form of ATP, AMPPNP. The crystal structure reveals that the HtpG_Mtb_ dimer adopts a conformationally silent structure, that precludes the dimerisation of the chaperone catalytic domains needed for ATP hydrolysis. Also, binding studies show that HtpG_Mtb_ directly interacts with TLR4 with a nanomolar affinity, and that this interaction allows HtpG_Mtb_ dimer to engage two TLR4 molecules. This finding suggests that activation of TLR4 by HtpG_Mtb_ is due to its ability to induce intra-cellular receptor dimerisation in an LPS-like mode.

## Introduction

Tuberculosis (TB) is caused by the ancient bacterium *M. tuberculosis*. It primarily infects alveolar macrophages and can produce lung cavitation, due to collagen degradation resulting from a strong immune response (Delogu *et al*, 2013; Squeglia *et al*, 2018b). However, some of the most devastating clinical consequences of TB arise from the ability of *M. tuberculosis* to spread from the lungs to other organs. Extra-pulmonary TB constitutes 15–20% of all TB cases and can be challenging to diagnose due to its varied presentations and locations (Baykan *et al*, 2022). According to WHO, 10.8 million people were infected with TB globally in 2023, causing about 1.25 million deaths. In addition, one-fourth of the world’s population is infected with a dormant form of *M. tuberculosis* (Gutti *et al*, 2019).

Dormancy of *M. tuberculosis* is its ace in the hole. Indeed, infected cells are sequestered in the TB granuloma, a niche where *M. tuberculosis* can persist in a dormant state for decades (Dutta & Karakousis, 2014; Romano *et al*, 2023). During dormancy, *M. tuberculosis* cells are not able to replicate and to be detected in biological samples, but this non-culturable state is transient (Mukamolova *et al*, 2005). The reactivation from dormancy occurs in about 10% of infected individuals, in a complex phenomenon that has been associated with the peptidoglycan degrading action of a set of proteins, denominated resuscitation promoting factors (Kana & Mizrahi, 2010; Ruggiero *et al*, 2009; Nikitushkin *et al*, 2015; Rosser *et al*, 2017; Squeglia *et al*, 2018a).

Dormancy is also a radical case of stressful environment, as it establishes conditions that would normally be harmful to cellular proteins, which typically undergo denaturation and aggregation (Batyrshina & Schwartz, 2020; Schramm *et al*, 2020; Vaubourgeix *et al*, 2015). However, *M. tuberculosis* is able to survive under the chemically stressful conditions including acidic pH, reactive oxygen, nitrogen species, antibiotics (Ehrt & Schnappinger, 2009; Baker *et al*, 2019; Harnagel *et al*, 2021; Levitte *et al*, 2016). Proteomic analyses has revealed the high abundance of proteins with a stabilising and protective effect in different dormancy models (Albrethsen *et al*, 2013; Schubert *et al*, 2015). These include chaperones HtpG_Mtb_ (Rv2299), DnaK (Rv0350), DnaJ1 (Rv0352), GroEL2 (Rv0440), GroES (Rv3418), GroEL1 (Rv3417) (Vaubourgeix *et al*, 2015; Kwiatkowska *et al*, 2008; Maisonneuve *et al*, 2008; Navarro Llorens *et al*, 2010; Starck *et al*, 2004; Trutneva *et al*, 2020; Albrethsen *et al*, 2013; Shleeva *et al*, 2011; Fay & Glickman, 2014). A finely tuned expression of this chaperone team is critical for *M. tuberculosis* response to conditions of stress (Harnagel *et al*, 2021). While DnaK is essential for cell growth and protein folding in *M. tuberculosis* (Fay & Glickman, 2014), HtpG_Mtb_ and ClpB are not essential (Vaubourgeix *et al*, 2015; Lopez Quezada *et al*, 2020). However, cells lacking both chaperones are oversensitive to host induced stresses (Harnagel *et al*, 2021). Consistently, HtpG_Mtb_ exhibits chaperonin activity in coordination with the DnaK/DnaJ/GrpE chaperone system, via direct association with DnaJ2 (Mangla *et al*, 2023; Berisio *et al*, 2024). Also, expression of DnaJ1, DnaJ2, ClpX, and ClpC1 increases in a ΔHtpG mutant strain of *M. tuberculosis* (Mangla *et al*, 2023).

HtpG_Mtb_ belongs to the highly conserved Hsp90 family of protein chaperones. Its homolog from *E. coli* (46% sequence identity) is involved in remodeling and activation of heat-inactivated proteins, in concert with DnaK (Genest *et al*, 2011). Importantly, HtpG_Mtb_ is highly conserved in pathogenic mycobacteria, like *M. leprae*, but not encoded in avirulent species, like *M. smegmatis*. A large number of structural studies of *M. tuberculosis* chaperones has been conducted in the last few years (Moreira *et al*, 2020; Yin *et al*, 2021; Xiao *et al*, 2024; Guillet *et al*, 2019). However, no experimental structural data are available for the chaperone HtpG_Mtb_, likely due to its strongly dynamic nature, typical of Hsp enzymes (Mader *et al*, 2020). Indeed, it was previously shown that the catalytic activity couples to conformational changes in these molecular chaperones (Mader *et al*, 2020; Moreira *et al*, 2020). All chaperones belonging to the Hsp90 family are homodimers with each protomer consisting of three major domains: the N-terminal (N), middle (M), and C-terminal (C) domains. The N domain is the site of ATP binding, the M domain is involved in ATP hydrolysis and client protein recognition, whereas the C domain is the site of constitutive dimerisation (Prodromou, 2016).

Besides being important for protein folding, HtpG_Mtb_ was shown to induce a strong activation of dendritic cells (DCs), and to enhance the protective immune response when fused to the known *M. tuberculosis* antigens ESAT6 (Mangla *et al*, 2023; Choi *et al*, 2017; Moreira *et al*, 2020; Ruggiero *et al*, 2022). In previous studies, we showed that, like other Hsp proteins, HtpG_Mtb_ is a dimeric nucleotide-binding protein and identified the key residues involved in the enzyme dimerisation, located on the C-terminal domain of the protein (Moreira *et al*, 2020; Ruggiero *et al*, 2022). We also found that the catalytic domains on the HtpG_Mtb_ dimer play no role in the elicitation of the immune response, which is mainly induced by the C-terminal and middle domains of the protein and experimentally validated these predictions in mice (Ruggiero *et al*, 2022). Also, we showed that the dimeric state of HtpG_Mtb_ favours ESAT6 dimerisation and hampers ESAT6 cytotoxicity (Moreira *et al*, 2020).

Many questions remain open on the structural determinants of HtpG_Mtb_ foldase activity, given the lack of structural data of HtpG_Mtb_. A key issue is the role of adenosine nucleotides in the clamping mechanism typical of Hsp proteins (Gu *et al*, 2025). To address this issue, we have determined the crystal structure of HtpG_Mtb_ in complex with AMPPNP, the non-hydrolysable derivative of ATP. Another open question is whether there is direct involvement of Toll Like Receptors in the DC maturation induced by HtpG_Mtb_ (Choi *et al*, 2017). We show here that HtpG_Mtb_ can bind TLR4 with binding affinity in the nanomolar range and that this interaction is mediated primarily by the C domain, assisted by the M domain. Together, our data support a complex clamping model, in which chaperone activation via N-terminal domain dimerisation requires not only ATP binding but also additional factors such as client proteins or co-chaperones. In addition, the observed ability of HtpG_Mtb_ to simultaneously engage two TLR4 receptors offers a mechanistic rationale for the TLR4-dependent activation of dendritic cells (Choi *et al*, 2017).

## Results

### Overall structure of HtpG_Mtb_ dimer in its ATP-bound state

Recombinant full length HtpG from *M. tuberculosis* (HtpG_Mtb_) was produced in E. coli and crystallised in complex with AMPPNP, the non-hydrolysable form of ATP. Crystallisation trials were performed using the Mosquito robotic workstation and then set manually using the hanging drop method. The best diffracting crystals were obtained using the hanging drop method and a precipitant solution composed of 20% (v/v) PEG 3,350 and 0.2 M MgCl_2_ and in the presence of AMPPNP/Mg^2+^ 4 mM. Structural determination was achieved using molecular replacement and the program Autorickshaw (Panjikar *et al*, 2005). All attempts using the structures of HtpG_Mtb_ homologues or the AlphaFold3.0 model as starting templates were unsuccessful. The structure solution could be obtained using the isolated middle and C-terminal domains from the AlphaFold3.0 model (Abramson *et al*, 2024), this indicating a different orientation of domains in the crystal structure, compared to the template models. After completing the building and refinement steps using the program Phenix (Adams *et al*, 2010) the final model contained amino 1242 acid residues, two AMPPNP molecules and two Mg^2+^ ions. Refinement statistics related to the final model are shown in Table 1.

**Table 1.** Crystallographic data processing and refinement statistics. Values in parentheses are for the highest resolution shell (3.4 – 3.3 Å). ^†^R_merge_= Σ hΣi |*I*(*h*,*i*)-<*I*(*h*)>|/ Σ hΣi *I*(*h*,*i*), where *I*(*h*,*i)* is the intensity of the *i*th measurement of reflection *h* and <*I*(*h*)> is the mean value of the intensity of reflection *h*.

| **Data Collection** |  |
| --- | --- |
| **Space group** | P 2<sub>1</sub>2<sub>1</sub>2 |
| **Unit-cell param. A,b,c (Å); $\alpha,\beta,\gamma$ (°)** | 173, 83.1, 98.5; 90°90° |
| **Resolution range (Å)** | 75.3-3.3 |
| **Average mosaicity** | 0.2 |
| **Multiplicity** | 10.2 (10.5) |
| **Total reflections** | 222368 |
| **Unique reflections** | 21831 |
| **Completeness (%)** | 99.4 (99.8) |
| $^{\dagger}R_{\text{merge}}$ | 0.11 (0.45) |
| **Average I/$\sigma$(I)** | 9.0 (0.6) |
| **Refinement** |  |
| **Rwork/Rfree (%)** | 26.1 (31.0) |
| **No. of residues** | 1242 |
| **R.m.s. deviations:** |  |
| **bond lengths (Å)** | 0.02 |
| **bond angles (°)** | 0.6 |

The crystal structure of HtpG_Mtb_-AMPPNP complex adopts a dimeric arrangement, consistent with our previous studies and with the functional role of HtpG_Mtb_ as a molecular chaperone (Figure 1) (Ruggiero *et al*, 2022; Moreira *et al*, 2020). However, the organisation of N, M and C domains of the structure unveiled unexpected structural details. Indeed, different than expected for an active ATP-bound state (Huck *et al*, 2017), HtpG_Mtb_ complex with AMPPNP does not adopt a closed N-terminally dimerised state, but it arranges as a V-shaped homodimer, with the dimerising C-terminal domains as the vertex (Figure 1A). It was previously shown that the engagement of the N-terminal catalytic domains in dimer formation to form the activated “close” conformation, is needed to allow the catalytic loop residue Arg380 (Arg 353 in HtpG_Mtb_) of the middle domain to access and complete the ATP-binding site (Ali *et al*, 2006) (Figure S1). In HtpG_Mtb_-AMPPNP structure, only a few inter-chain interactions are mediated by residues of N domains, including hydrogen bonds between Ser150 side chain and the main chain N of Lys55 and Asp56 of the adjacent chain, and a salt bridge between Arg88 (chain B) and the side chain of Glu354 (chain A)(Figure 1B). Instead, the dimerisation interface covers 1684.0 Å^2^ (as computed by PISA), only 5% of the entire solvent accessible surface area (32867.3 Å^2^) and is almost solely formed by the C-terminal domains of the enzyme (Figure 1).

**Figure 1.**
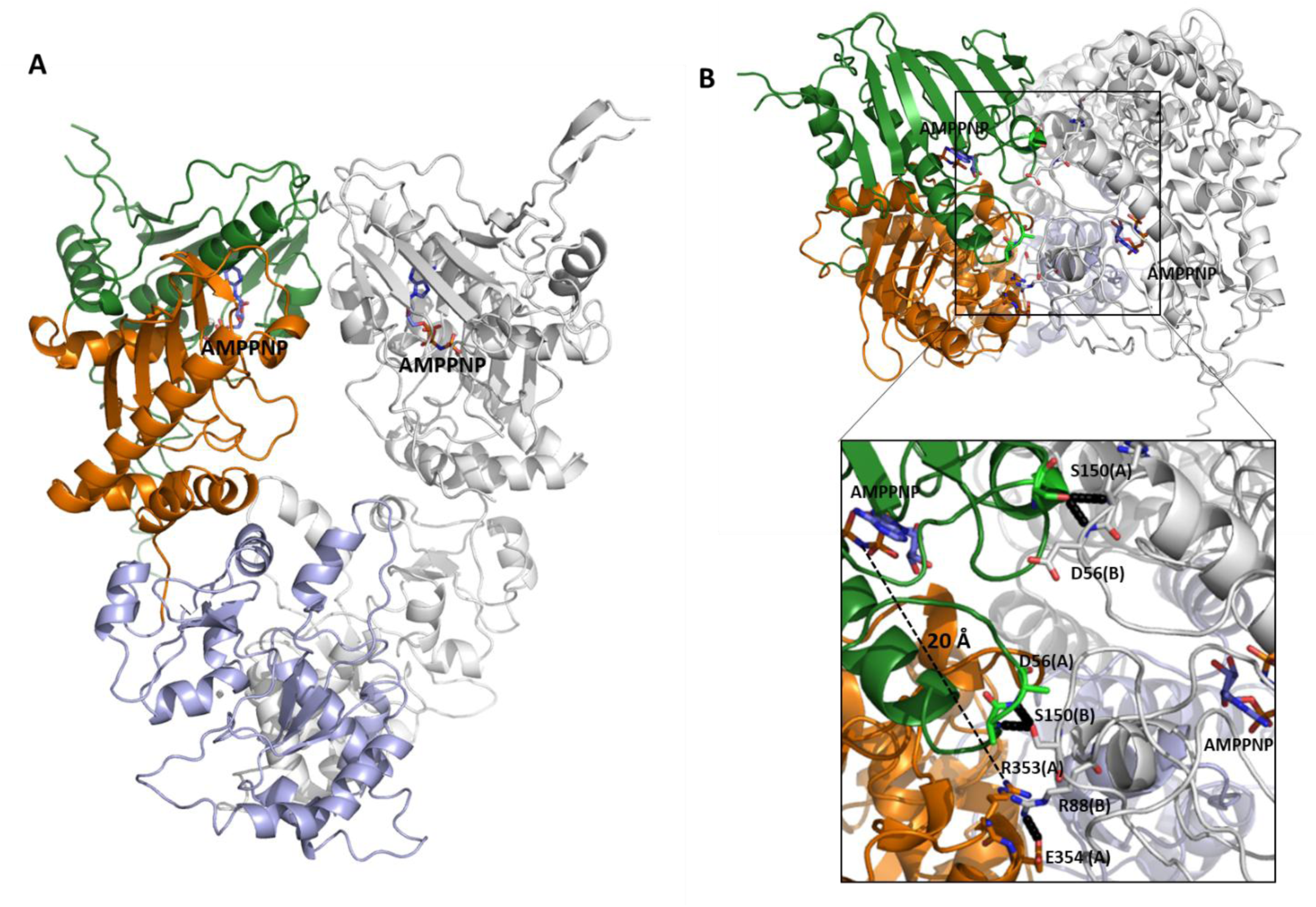
Cartoon representation of HtpG_Mtb_ crystal structure in side view (left) and top view (right). In chain A, the N-terminal domain is drawn in green, the M domain in orange and the C domain in light blue. The B chain is drawn in grey, for clarity. AMPPNP molecules bound to both chains A and B are drawn in stick. In panel B, the inset shows a close up of residues, drawn in stick, involved in interactions mediated by the N-terminal domain. The distance between Arg353 side chain and the imido linkage of the phosphoamidate bond between the β and γ phosphates of AMPPNP (modified version of the phosphate bond) is represented by a dashed line.

Alignment of the HtpG_Mtb_-AMPPNP with known structures, performed using DALI, identifies the apo structure of *E. coli* HtpG in the open state as the most similar (PDB code 2ioq, Z score 39.3, RMSD 3.7 Å). Superposition of the protomers shows that the N, M and C domains of these structures are oriented similarly (Figure 2). However, a closer angle between the two protomers characterises HtpG_Mtb_-AMPPNP complex compared to the unliganded conformation of HtpG_Mtb_ from *E. coli* (Figure 2)(PDB code 2ioq). In this conformation, here denominated V-shape, predicted catalytic residues (Glu35) from the two protomers of HtpG_Mtb_ are located 45.5 Å from each other, whereas the equivalent residues in *E. coli* HtpG (Glu34) are much further, 85.3 Å away (Figure 2). Importantly, the catalytic domains in HtpG_Mtb_-AMPPNP complex are oriented as in *E. coli* HtpG, whereby they point in directions that are diametrically opposed to those required for dimerisation (Figure 2). As previously explained (Figure 1B), in the observed V-shape conformation, Arg353 of the middle domain is more than 20 Å away from the γ-phosphate group of the AMPPNP molecule, too far to assist the ATP hydrolysis reaction.

**Figure 2.**
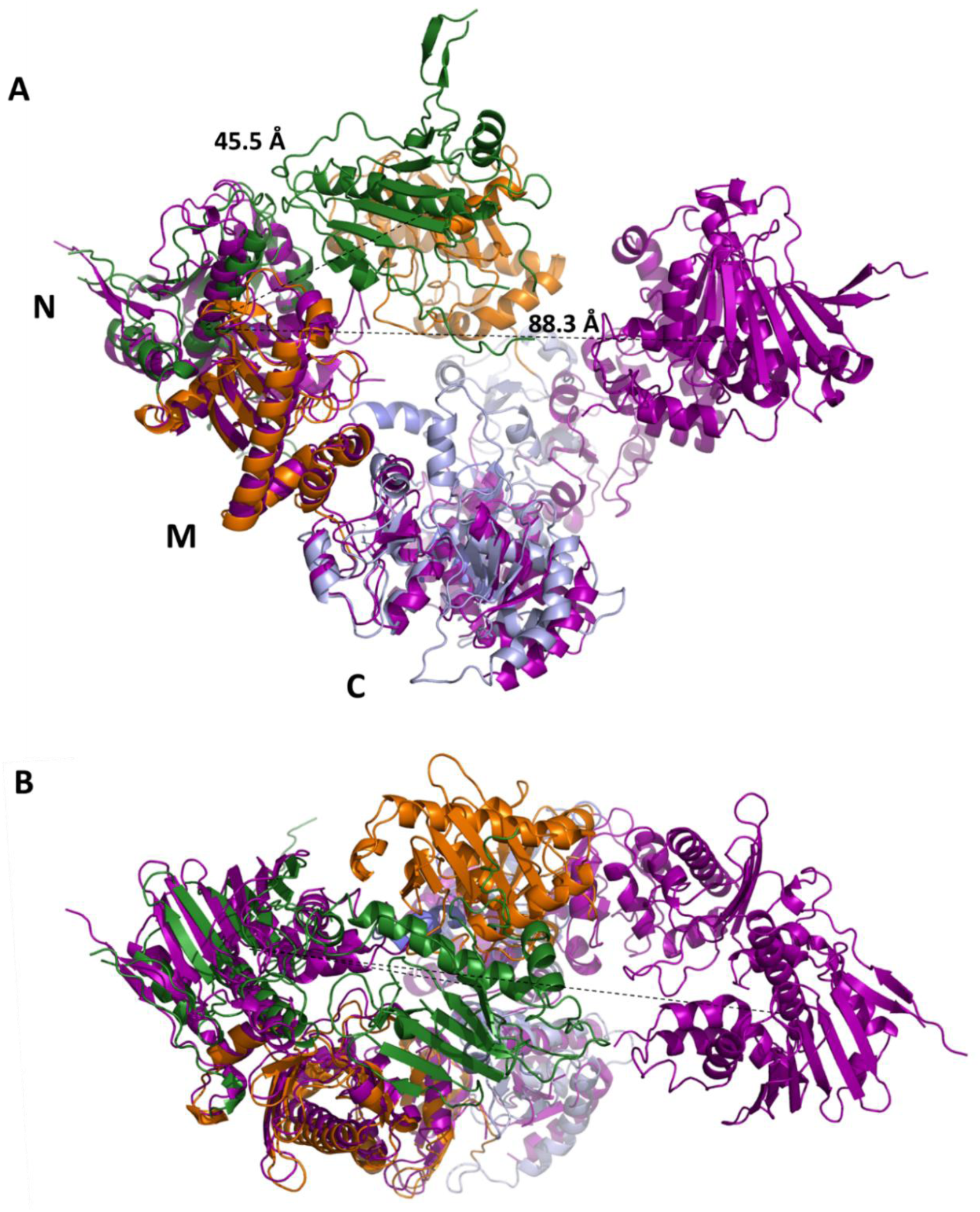
Superposition of the structure of HtpG_Mtb_-AMPPNP complex with that of HtpG from *E. coli* (prune, PDB 2ioq). In HtpG_Mtb_-AMPPNP structure, N-terminal domain is drawn in green, the M domain in orange and the C domain in light blue. Lines represent the distances between the predicted catalytic E35 (E34 in HtpG from *E. coli*) of the two protomers in each structure.

Superposition of HtpG_Mtb_ to the “close dimer” conformation of Hsp90 from *Saccharomyces cerevisiae* (PDB code 2cg9) using DALI (Holm, 2022) (Z score 21.0) reveals large conformational difference both in the organisation of domains in each protomer and in the assembly of protomers (Figure 3A)(Ali *et al*, 2006). Analogous conformational change is observed with the close conformation of the eukaryotic endoplasmic reticulum Hsp90 from *canis lupis familiaris* GRP94 (PDB code 5uls)(Huck *et al*, 2017). Overall, HtpG_Mtb_ appears less elongated, compared to the close state (Figure 3A). When A chains are superposed, the N domain of the AMPPNP-bound HtpG_Mtb_ is twisted, with a rotation about the N-M interdomain junction by ∼90 degrees, compared to the typical “closed state” (Figure 3B). This conformational variation does not allow in HtpG_Mtb_ the dimerisation of the N domains observed in yeast Hsp90. Instead, N-terminal ends point in directions opposite to those necessary for the domain swapping observed in the closed state (Figure 3).

**Figure 3.**
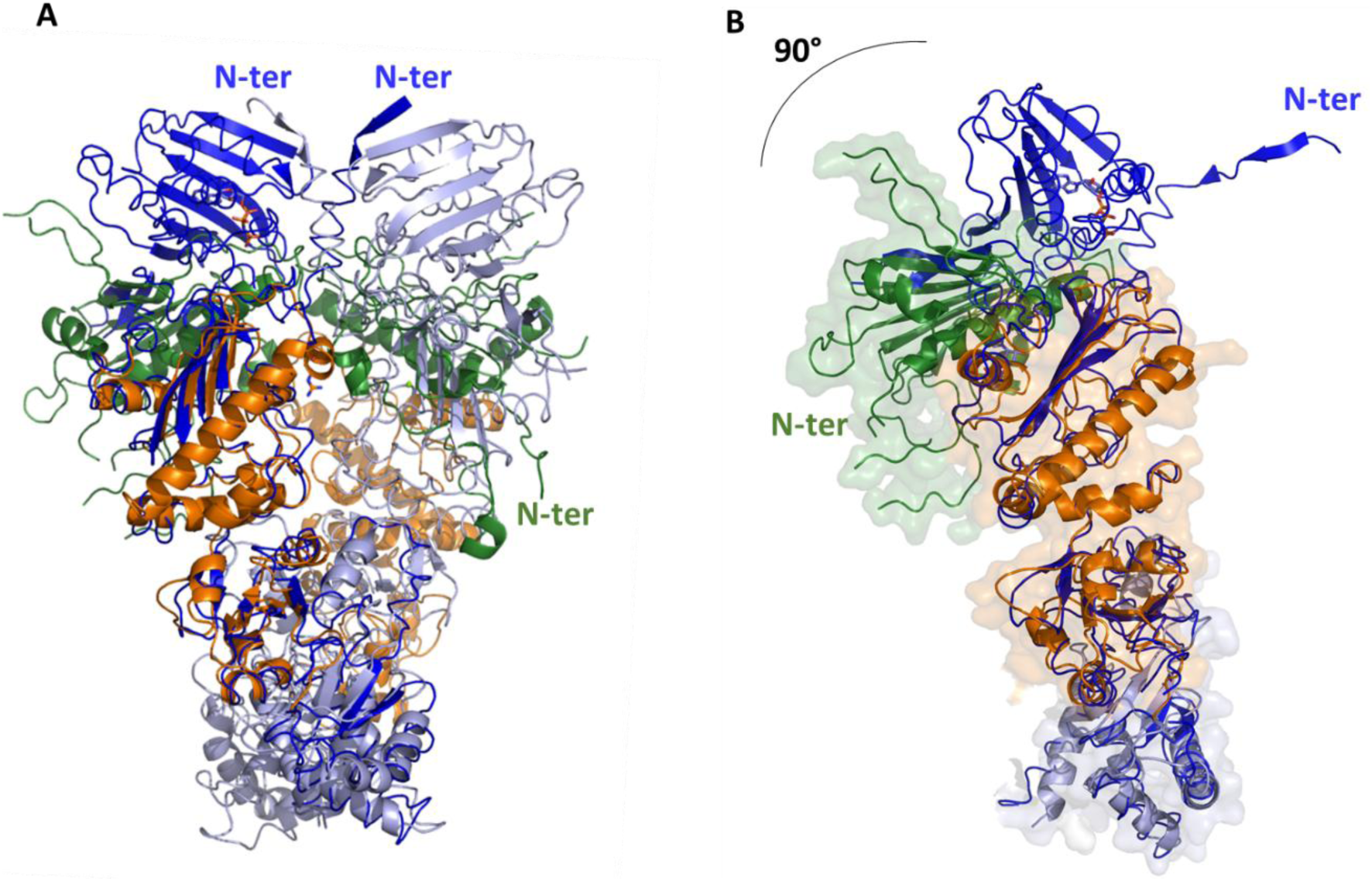
Superposition of the structure of HtpG_Mtb_-AMPPNP complex with that of Hsp90 from *Saccharomyces cerevisiae* in the closed state. (A) Cartoon representation of HtpG_Mtb_-AMPPNP complex structure (N-terminal domain is drawn in green, the M domain in orange and the C domain in light blue) and Hsp90 from *Saccharomyces cerevisiae* (PDB code 5cg9, blue). N-terminal ends for the two enzymes are labelled with the same color of the catalytic domains. In HtpG_Mtb_-AMPPNP, one N-ter is not visible because behind the plane of the drawing. (B) Cartoon and surface representation of protomer A of HtpG_Mtb_-AMPPNP, superposed to protomer A of Hsp90 from *Saccharomyces cerevisiae* in cartoon, for sake of clarity. This orientation (60° rotated compared to panel A) was chosen to highlight the 90° rotation of the catalytic domain in HtpG_Mtb_-AMPPNP, compared to yeast Hsp90 (PDB code 5cg9).

### The crystal structure of HptG_Mtb_ AMPPNP complex reveals a strikingly different conformation compared to AlphaFold3 prediction

The impressive success of AlphaFold3.0 in producing highly accurate structural models has led to a surge in studies relying exclusively on its analyses (Abramson *et al*, 2024). Nevertheless, it is crucial to recognise that AlphaFold3 approach is grounded in previously determined experimental structures from the PDB database, rather than in biophysical principles and its predictions may be influenced or biased by the existing structural data. In the case presented here, AlphaFold 3.0 predicts a structural model with high confidence (average plDDT 85.0, ipTM 0.78, pTM 0.81). However, this conformer is strikingly different from that observed in the crystal structure, with RMSD computed on Cα atoms of the of each protomer of 5.0 Å. Indeed, the Alphafold3.0 model strongly resembles the close swapped dimer conformation, with a 90° rotated catalytic domain, when M domains are superposed (Figure S2). Consistently, molecular replacement using the full AlphaFold3.0 model was unsuccessful (Abramson *et al*, 2024). These findings serve as a cautionary reminder that, although the structural models produced by AlphaFold3.0 are impressive, they must be validated through thorough experimental investigation. On the other hand, also this discrepancy witnesses the intrinsically high mobility of domains in large Hsp structures, a finding that makes predictions less reliable. High interdomain mobility is indeed a fundamental characteristics for the biological function of Hsp molecules, able to completely change shape in the different functional states, to either engage new proteins to fold or to release already folded proteins.

### HptG_Mtb_ substrate binding and ATPase activity

The analysis of electron density maps clearly allowed the identification of AMPPNP molecules bound to the catalytic site of each chain (Figure 4A). AMPPNP is kept in its binding site through hydrogen bonds of its β- and γ-phosphates with Asp35 and Asn39, and several hydrophobic interactions with Met86, Leu94, Phe130 (Figure 4A). Superposition of the catalytic domain of HtpG_Mtb_-AMPNP complex with that of the closed active conformation of Hsp90 from *Saccharomyces cerevisiae* (Ali *et al*, 2006) provides interesting clues. In previous MD analyses, we showed that the main conformational changes between the different functional states of HtpG_Mtb_ are due to the interplay of inter-domain motions and the conformational variation of a region, denoted as lid (residues 99-130), to allow nucleotide binding (Moreira *et al*, 2020). In the closed conformation (PDB code 2cg9), the catalytic pocket is completely closed by the lid and the ATP γ-phosphate is locked in the catalytic pocket by the side chain of Arg380 (Arg353 in *M. tuberculosis*), belonging to the M domain (Figure 4B, Figure S1). Despite the conservation of the nucleotide binding pocket in the HtpG_Mtb_-AMPPNP crystal structure, the lid region is in a fully open conformation and exposes the nucleotide binding pocket (Figure 4B). As previously mentioned and shown in Figure 1B, different than in the closed state (Figure 4B), Arg353 of the M domain does not participate to AMPPNP binding, but it is displaced more than 20 Å away (Figure 1B), suggesting that the structure of HtpG_Mtb_-AMPPNP complex is trapped in a catalytically incompetent state. These structural features prompted us to check whether the protein is at all endowed with any ATPase activity and to identify potential catalytic residues.

**Figure 4.**
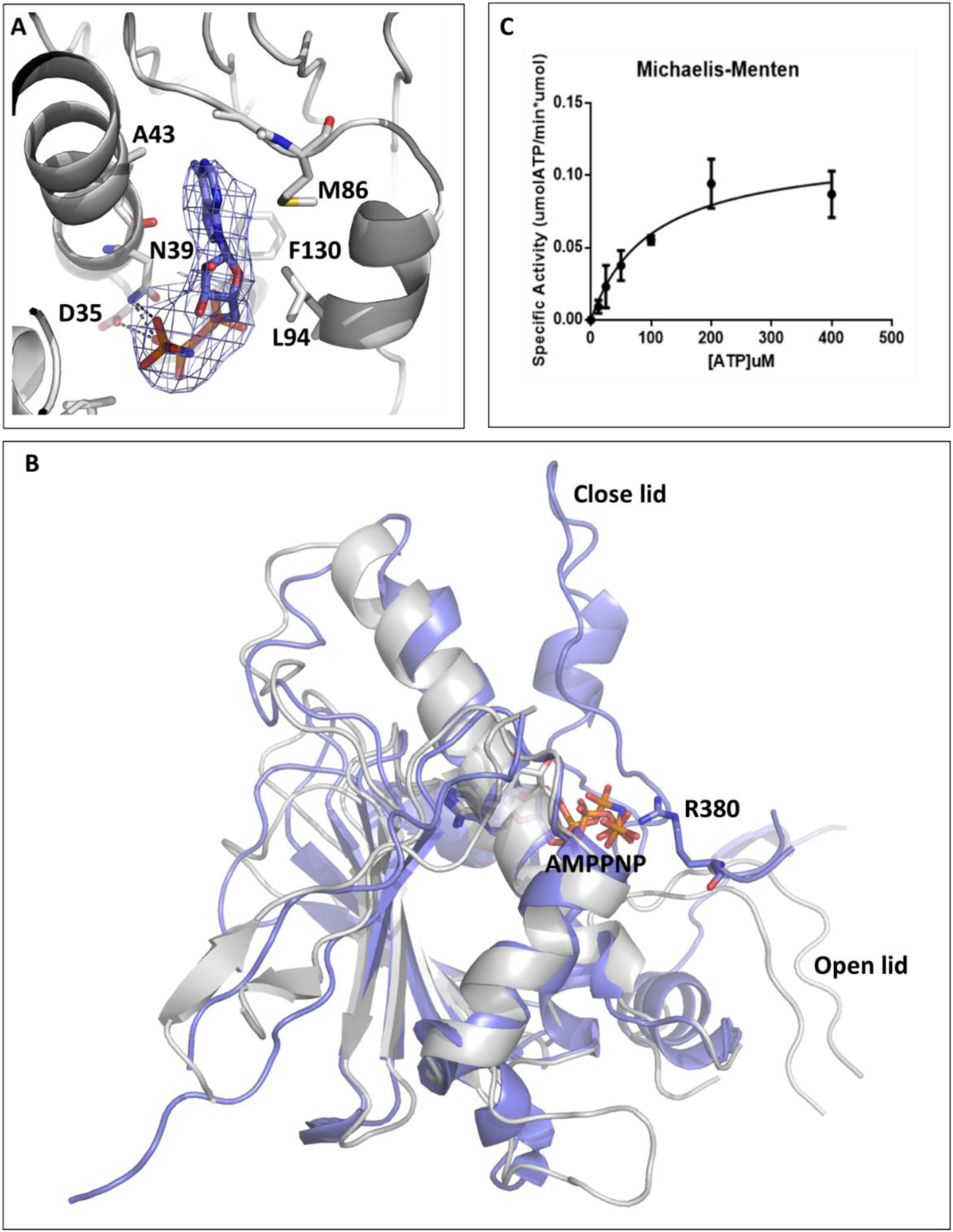
Effect of AMPPNP binding on the catalytic site. (A) Stick representation of HtpG_Mtb_ catalytic site (chain A). The omit (Fo-Fc) electron density map, contoured around AMPPNP at 2σ, is shown in blue. (B) Conformational differences of HtpG_Mtb_ catalytic domain (grey) compared to the closed state of Hsp90 from *Saccharomyces cerevisiae* (blue, PDB code 2cg9). Completely different lid conformations are observed, from fully close in Hsp90 to an open lid in HtpG_Mtb_ structure. (C) ATPase activity of HtpG_Mtb_. Reactions were performed with 5 μM of protein using different concentrations of ATP spanning from 1600 μM to 50 μM in the presence of 5 mM MgCl_2_ in 50 mM TrisHCl (pH 7.5). Kinetic parameters such as Km and Vmax were determined using GraphPad Prism. All the reactions were performed in triplicate, and the mean ± standard deviation (SD) values are shown.

Using a PiPer Phosphate Assay (Invitrogen), we found that kinetic parameters are indicative of low ATPase activity, with a Kcat and Km measured at 37°C and pH 7.5 of 0.12 ± 0.013 min^-1^ and 100 ± 28 mM, respectively, and catalytic efficiency (Kcat/Km) of 1.2 10^-3^ min^-1^ mM^-1^ (Figure 4C). Based on the typical reaction mechanism of ATPases (Mader *et al*, 2020), we predicted that (among AMPPNP contacting residues, Figure 4A), Glu35 serves as a key catalytic residue, to properly position and polarise a lytic water molecule for ATP hydrolysis (Mader *et al*, 2020). To experimentally confirm Glu35 as a key catalytic residue, we mutated it to alanine and measured the catalytic efficiency of this mutant. Results show that mutation of Glu35 leads to a complete loss of HtpG_Mtb_ ATP-degrading activity. The poor ATPase activity of HtpG_Mtb_ suggests that, unlike other members of the GHKL family, ATP binding alone does not drive HtpG_Mtb_ into a hydrolytically productive conformation, but another event is needed to induce the catalytically competent state. Also, the non-swapped structure of AMPPNP-bound HtpG_Mtb_ explains the poor discrimination we previously measured for binding of ADP and AMPPNP (Ruggiero *et al*, 2022).

### HtpG_Mtb_ directly binds the toll like receptors TLR4 with nanomolar affinity

Toll-like receptors (TLRs) play critical roles in the innate recognition of *M. tuberculosis* by host immune cells. HtpG^Mtb^ was shown to activate TLR4 receptor, although its activation mechanism is hitherto unknown (Choi *et al*, 2017). We used Surface Plasmon Resonance (SPR) to check whether HtpG_Mtb_ is able to directly bind TLR4 and provide a detailed description of the interaction event. Real-time binding assays were performed to assess binding kinetics and affinity between HtpG_Mtb_ and its shorter variants toward TLR4 and TLR4/MD2, using Biacore T200 (Cytiva, Uppsala, Sweden). In parallel experiments, either TLR4 or TLR4/MD2 were stably captured at the surface of the CM5 sensor chip, using standard amine-coupling protocols (see methods). Following immobilisations, protein solutions were injected at various concentrations (from 0.002 to 1 µM), using a flow rate of 30 μL/min for 120 s (association phase), and then the buffer alone for 300 s (dissociation phase). Equilibrium dissociation constant (K_D_) values were derived from the ratio between kinetic dissociation (kd) and association (ka) constants, obtained by fitting data from all injections at different concentrations of each compound, using the simple 1:1 Langmuir binding fit model of the BIAevaluation software (version 2.0.2). SPR analysis shows that full length HtpG_Mtb_ binds to either TLR4 or TLR4/MD2 with elevated affinity, with K_D_ values in the low nanomolar range (Table 2, Figure 5). With the aim to identify relevant regions of HtpG_Mtb_ involved in TLR4 binding, we recombinantly produced more variants of the enzyme including, beside the full-length protein, the sole C-terminal domain (HtpG_C_, residues 420-647), the middle M domain (HtpG_M_, residues 248-420), and both M and C domains (HtpG_MC_, residues 248-647)(Figure 5A). SPR analysis shows the strongest binding for HtpG_MC_ to TLR4 and TLR4/MD2 species, with K_D_ values comparable with those of the full-length protein. This result is consistent with our previous findings that the catalytic N-terminal domain does not play any role in HtpG-induced elicitation of the immune response (40). Further analyses show that HtpG_C_ binds to the TLR4 receptor with lower affinity compared to HtpG_MC_, albeit still in the nanomolar range (Table 2). On the other hand, a K_D_ value in the low micromolar range characterises TLR4 binding of HtpG_M_ (Table 2, Figure 5). These data consistently show that binding to TLR4 involves mainly the C-terminal domain and part of the M domain of HtpG_Mtb_.

**Figure 5.**
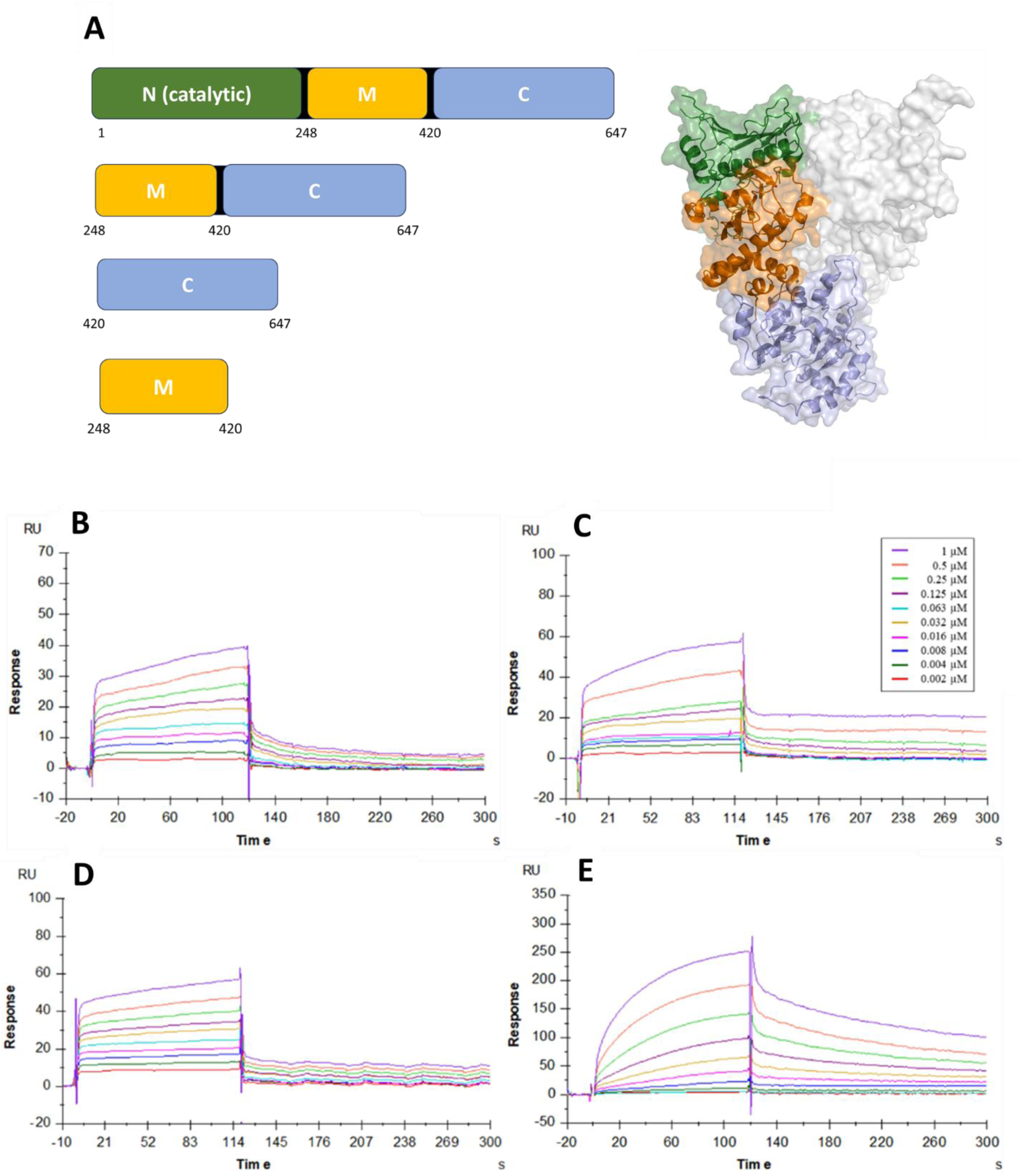
Surface plasmon resonance (SPR) analysis. (A) Scheme of the assayed recombinant HtpG_Mtb_ variants (left). For clarity, a surface representation of HtpG_Mtb_ structure is reported (right) with the same colour code. (B) Sensorgrams to evaluate the binding of HtpG_Mtb_, HtpG_C_ (C), HtpG_MC_ (D) and HtpG_M_ (E) with TLR4/MD2.

**Table 2:** Dissociation constant (K_D_) values (µM), determined by SPR, for the interaction with TLR receptors with HtpG_Mtb_ and its shorter variants

|  | **$K_D$ values ($\mu\text{M}$)** |  |  |  |
| --- | --- | --- | --- | --- |
|  | HtpG | HtpG<sub>MC</sub> | HtpG<sub>C</sub> | HtpG<sub>M</sub> |
| TLR4 | 0.04±0.02 | 0.028±0.008 | 0.44±0.06 | 5.8±0.3 |
| TLR4/MD2 | 0.02±0.01 | 0.044±0.003 | 0.14±0.04 | 4.45±0.02 |

### HtpG_Mtb_ interaction prompts TLR4 activation through LPS-like dimerisation

We have previously shown that HtpG_Mtb_ activates dendritic cells (DCs), leading to cytokine production and upregulation of co-stimulatory molecules, in a TLR4-dependent mechanism. Indeed, blocking TLR4 or using TLR4-deficient cells significantly reduced the immune response (Choi *et al*, 2017). Despite the importance of this event in triggering an NF-κB-mediated immune cascade, the exact structural mechanism of TLR4 activation is hitherto unknown. A well-established mechanism of TLR4 activation is triggered by the binding of lipopolysaccharide (LPS), a key component of the outer membrane of Gram-negative bacteria. The process begins when LPS, particularly its lipid A moiety, is recognised by a complex of accessory proteins, including LPS-binding protein (LBP), CD14, and MD2. CD14 facilitates the transfer of LPS to the TLR4-MD2 complex located on the surface of immune cells such as macrophages and dendritic cells. Upon LPS binding, TLR4 undergoes homodimerisation, which triggers intracellular signaling cascades. These pathways activate transcription factors such as NF-κB and IRF3, leading to the production of proinflammatory cytokines and type I interferons (Luo *et al*, 2025). Outcome of TLR4 activation is the triggering of two possible signalling cascades: the MyD88-dependent pathway (Ve *et al*, 2017) and the TRIF-dependent pathway (Manik *et al*, 2025), which drive NF-κB-mediated pro-inflammatory cytokine production (similar to HtpG_Mtb_) (Choi *et al*, 2017) and IRF3-induced type I interferon responses, respectively.

Building on this mechanism, we investigated whether HtpG_Mtb_ can induce TLR4 dimerisation in a manner analogous to LPS. Using SPR, we stably immobilised HtpG_Mtb_ on the surface of the CM5 sensor chip with standard amine-coupling protocols (see methods). Following immobilisation, TLR4/MD2 solutions were injected at concentrations from 0.002 to 1µM, using a flow rate of 30 μL/min for 120 s (association phase), and then the buffer alone for 300 s (dissociation phase). Stoichiometry of binding was evaluated using eqs 1:

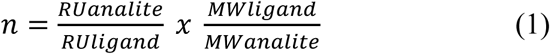

The equilibrium dissociation constant (K_D_) values, evaluated using a sequential binding mode, revealed two distinct binding events with KD1=0.46±0.02 µM and KD2 = 0.012±0.001 µM. These values show the presence two TLR4/MD2 binding sites on the HtpG_Mtb_ dimeric protein, with the interaction occurring in two consecutive steps characterised by distinct kinetic parameters (Figure 6). Consistently, the apparent stoichiometry calculated from the SPR response indicates that, on average, two molecules of TLR4/MD2 bind per immobilised HtpG_Mtb_ dimeric molecule, yielding a TLR4/MD2:HtpG_Mtb_ stoichiometry of 1.8±0.1. As reported in Table 2, each TLR4 monomer specifically interacts with a region of HtpG_Mtb_ situated between the C-terminal and middle domains, highlighting a structurally coordinated mode of receptor engagement. These findings point to a TLR4 association mechanism in which each TLR4 monomer is engaged by a binding interface on the HtpG_Mtb_ dimer. In this event, HtpGMtb acts as a crosslinking agent to favour the dimerisation of intracellular TLR4 receptors, needed for as a starting event for the trigger of the NF-κB-mediated cytokine cascade.

**Figure 6.**
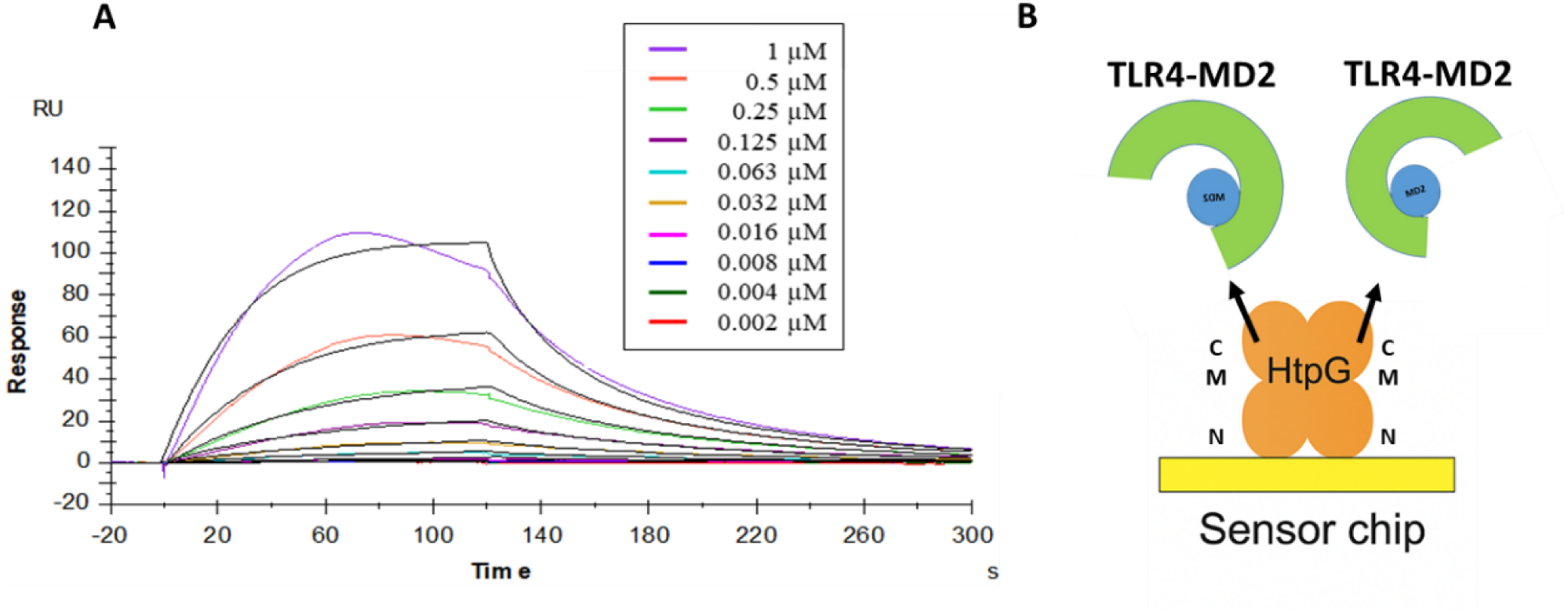
(A) Surface plasmon resonance (SPR) analysis. Sensorgrams to evaluate the kinetic affinity of TLR4/MD2 with HtpG_Mtb_. (B) A sketch exemplifying TLR4-MD2 recruitment by an HtpGMtb dimer on the sensor chip.

## Discussion

HtpG_Mtb_ is part of a complex chaperone network, including the chaperone DnaK and its co-factors DnaJ1/J2, the release factor GrpE and the disaggreases ClpB (Vaubourgeix *et al*, 2015; Yin *et al*, 2021; Lupoli *et al*, 2016; Fay & Glickman, 2014). The molecular mechanism by which this chaperone works and its role in innate immunity activation is still unknown, due to the lack of structural data. Here, we report the first crystal structure of HtpG_Mtb_, in complex with AMPPNP along with biophysical data that prove strong direct interaction with the host receptor TLR4.

ATP hydrolysis is closely associated with the function of all members of the Hsp family, and more models of hsp90 action propose that ATP binding alone drives the chaperone into a hydrolytically productive conformation (Pearl & Prodromou, 2006). The results presented here show that the binding of AMPPNP to HtpG_Mtb_ does not result in the “close” conformation observed for human (Prodromou, 2000) and yeast Hsp90 (Ali *et al*, 2006). In the reported “close” conformations (Prodromou, 2000; Ali *et al*, 2006), ATP binding induces a major conformational change where the two catalytic N-terminal domains of the dimer come together through domain swapping. Upon closure, a conserved arginine of the middle domain (Arg353 in *M. tuberculosis*) repositions to interact with the γ-phosphate of ATP; This creates a closed clamp around client proteins, essential for stabilising them during folding. Differently, HtpG_Mtb_ structure in complex with AMPPNP adopts a V-shaped conformation where catalytic domains do not undergo dimerisation. In this conformation, the Arg353 residues of the middle domains expected to bind ATP are located more than 20 Å away from the catalytic sites (Figure 1B), likely resulting in a catalytically incompetent state. Consistently, we show that HtpG_Mtb_ is endowed with poor ATP-ase activity (Figure 4C).

A similar conformation, never observed before for a bacterial or yeast Hsp, was found in the ATP complex of the mammalian GRP94, the endoplasmic reticulum paralog of human Hsp90 (Dollins *et al*, 2007). In the case of GRP94, this incompetent conformation was attributed to the lack to the pre-N region, located at the N-terminal end of the catalytic domain and absent in human Hsp90, that is essential for client maturation and important for the regulation of ATPase rates and dimer closure (Huck *et al*, 2017). Indeed, the nearly full-length protein including the pre-N domain (residues 48-72) was found to adopt the active “close” dimer (Huck *et al*, 2017). The behaviour of GRP94 was associated to a mechanistic difference that may distinguish mammalian hsp90s from their lower eukaryotic/bacterial homologs (Huck *et al*, 2017). Instead, our data show that HtpG_Mtb_ resembles in its behaviour the mammalian GRP94 more than yeast HtpG although it lacks the pre-N domain of GRP94 homologs (Huck *et al*, 2017).

Recent publications on the interactions of HtpG of *E. coli* with client proteins have highlighted that, despite the diversity of client proteins, their binding sites are located in close proximity (Qu *et al*, 2024b) and that client protein binding may act as an allosteric switch, that dynamically primes HtpG for elevated chaperone activity (Qu *et al*, 2024a). The poor ATPase activity and the V-shaped silent conformation that we observed for HtpG_Mtb_ (Figure 4B) suggest that the transition from the silent conformation to the “close” active conformation of HtpG_Mtb_ needs not only ATP binding but also additional factors such as the binding of client proteins and/or co-chaperones like DnaK and DnaJ2 (Berisio *et al*, 2024; Fay & Glickman, 2014).

Another important aspect we addressed here is the ability of HtpG_Mtb_ to induce maturation of DC in a TLR4-dependent manner (Choi *et al*, 2017). TLRs play an essential role in host immune responses to various invading pathogens, by recognising microbial membrane components to induce an inflammatory response and link innate and adaptive immunity by regulating the activation of antigen-presenting cells. The best known TLR4 activation mechanism is that induced by LPS, whose binding to TLR4-MD2 induces receptor dimerization and triggers two major signaling pathways: the MyD88-dependent pathway (Ve *et al*, 2017) and the TRIF-dependent pathway (Manik *et al*, 2025), leading to NF-κB–mediated pro-inflammatory cytokine production and IRF3-driven type I interferon responses, respectively. We show here that TLR4 recognises HtpG_Mtb_ at the molecular level through a strong binding, with nanomolar affinity, with the middle and C-terminal domains of HtpG_Mtb_, whereas the catalytic domains are not involved in TLR4 binding. Notably, our SPR data show that HtpG_Mtb_ dimers can simultaneously engage two TLR4 molecules. Based on this observation, it is tempting to surmise that HtpG_Mtb_ triggers an activation pathway analogous to the (myeloid differentiation primary response 88) MyD88-dependent pathway induced by LPS (Ve *et al*, 2017) (Figure 7). Similar to LPS, HtpG_Mtb_ facilitates the bridging of two TLR4 protomers, although receptor dimerisation by HtpG_Mtb_ is due to a direct crosslink of extracellular regions of TLR4. TLR4 dimerisation induced by HtpG_Mtb_ is essential for MyD88 recognition of the intracellular TIR domains of TLR4, and for initiating the entire signalling cascade (Ve *et al*, 2017). Without dimerisation, MyD88 cannot bind effectively, and the NF-κB/MAPK cascade is not triggered (Figure 7).

**Figure 7.**
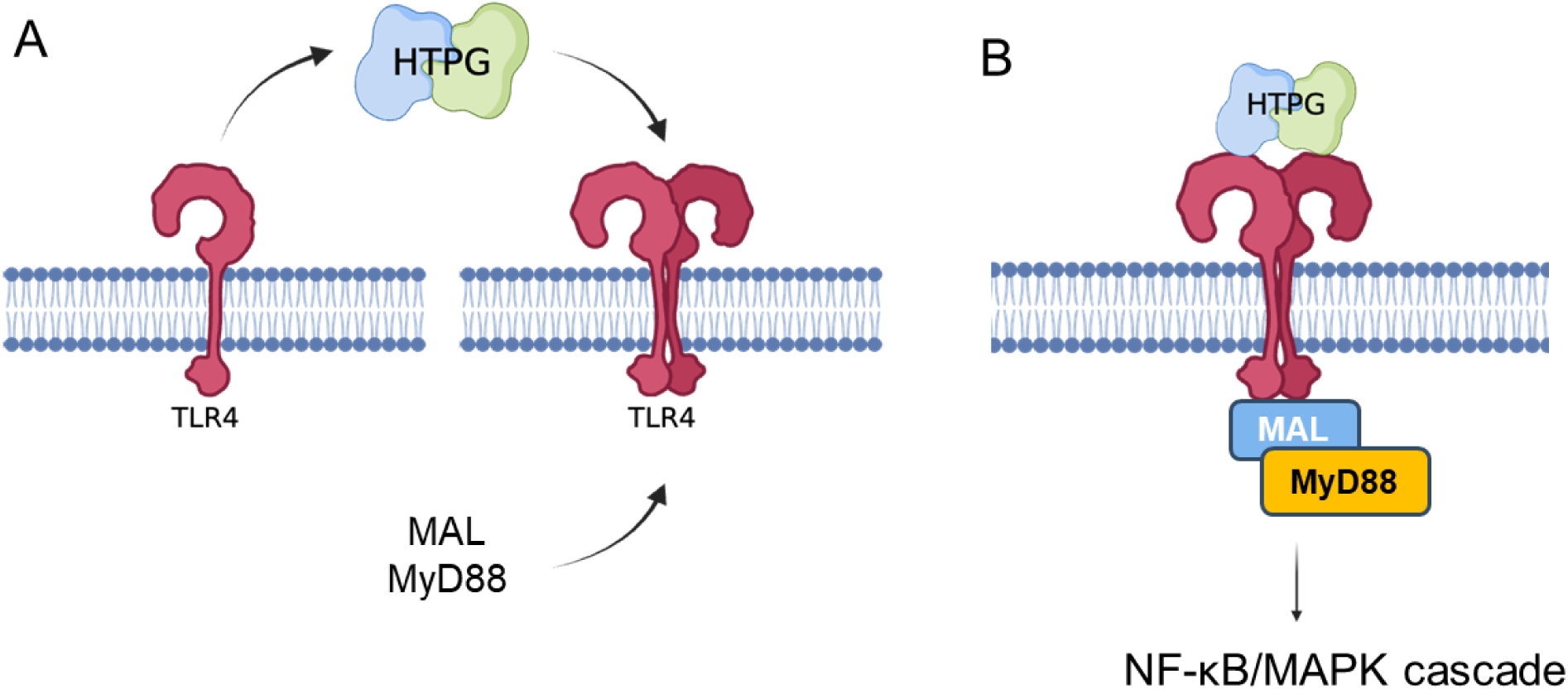
A mechanistic model for HtpG_Mtb_–induced TLR4 activation. (A) HtpG_Mtb_ is presented to the extracellular TLR4 monomers to induce TLR4 dimerisation. and consequent engagement of MAL and MyD88 adaptors. (B) The formation of a heterocomplex between HtpG_Mtb_, TLR4 and MAL/MyD88 adaptors through their interactions with intracellular dimeric TIR domains starts the NF-κB/MAPK cascade.

## Materials and Methods

### Protein Recombinant production

For the recombinant production of full-length HtpG_Mtb_ and of its truncated versions, we employed the expression vectors pET-22b(+) and pETM-13. Cloning procedures of designed genes encoding for MC- double domains, M- and C- single domain of HtpG_Mtb_ were based on PCR amplification using the Rv2299c gene as template and the primers reported in Table S1. Then, the resulting amplificated products were digested using NcoI and HindIII restriction enzymes and after purification were cloned into the pETM-13 expression vector using the T4 DNA Ligase (NEB Biolabs). The ATPase mutant (E35A) of HtpG_Mtb_ was generated using Q5 Site-Directed Mutagenesis Kit (NEB Biolabs) and the primers reported in Table S1. All the recombinant plasmids, propagated using *E. coli* Top10 competent cells, were recovered by using Qiagen mini-prep kit (Germany) and send to sequencing. Positive plasmids were used to transform BL21 (DE3) *E. coli* as expression strain, for a scale-up protein expression. Overnight precultures grown at 37°C, were inoculated in 1L LB-antibiotic flasks and the bacterial cells were grown at 37°C until an optical density at 600nm of 0.6 was reached to induce with 0.5 mM IPTG at 22°C.

The purification protocol was roughly similar in most cases. Briefly, the cell pellet was resuspended in 50 mL of lysis buffer (50mM Tris-HCl pH 7.8, 300mM NaCl, 5% (v/v) glycerol, 2 mM DTT), supplemented with a protease-inhibitor cocktail (Roche Diagnostics) and DNaseI and RNaseI. Sonication was set up for 10’ with a 21W ½ pulse and the cell lysate was cleared by centrifugation at 35,000xg for 45min at 4°C. The supernatant was loaded on Ni^2+^-NTA resin (Qiagen) equilibrated with a proper binding buffer. A series of washing steps were applied with small increasing of imidazole concentrations (10, 20, and 40mM), then the protein was eluted with 150-200 mM of imidazole. After elution, the fractions containing the protein were pooled and dialysed against a buffer of 50 mM Tris-HCl, 50mM NaCl, 5mM EDTA and 2mM DTT (pH 7.8) overnight at 4°C for salts removal. Indeed, further contaminants were removed by anionic exchange chromatography carried out on a 5 mL HiTrap Q HP column (Cytiva). The protein was eluted using a NaCl gradient ranging from 50 mM to 2 M. The collected fractions were analysed by SDS-PAGE and 1% (w/v) agarose gel electrophoresis to analyse the nucleic acid removal from protein solution. The fractions of interest were then collected and subjected to a second purification step on size exclusion chromatography. This step was carried out using a Superdex 200 increase 10/300 or Superdex 75 increase 10/300 column (Cityva) that were pre-equilibrated with running buffer containing 50 mM Tris-HCl, 150 mM NaCl, 5% (v/v) glycerol and 2 mM DTT (pH 7.8). The purified proteins eluted as a single peak, and the resulting fractions were analysed by SDS-PAGE to confirm protein purity and homogeneity. Finally, the fractions identified were pooled together, to concentrate the protein of interest for subsequent analyses in Amicon centrifuge concentrators (Millipore, Merck). Protein concentration (mg/mL) was determined using a Nanodrop OneC (ThermoFisher), using protein parameters, ε (M^-1^ cm^-1^) and MW (g mol^-1^) calculated by ProtParam tool on the ExPASy server (https://web.expasy.org/cgi-bin/protparam/protparam).

### Crystallisation and data collection of HtpG_Mtb_

First attempts for protein crystallisation were achieved by a high-throughput screening approach by using the sitting-drop vapor-diffusion technique at 293 K. To this aim the crystallisation space was explored using commercially available sparse-matrix precipitant solutions from Hampton Research kits (Crystal Screen I and II, INDEX, PEG-Ion USA). For crystallisation trials, freshly purified HtpG_Mtb_ in a concentration range from 5 to 7 mg/mL, was mixed with AMP-PNP/Mg^2+^ in a 1:10 molar ratio. Crystal growth was then manually optimised using the hanging-drop in 24-well plate where both protein and precipitant concentration were rationally varied.

To prepare the crystals for diffraction, they were frozen in liquid nitrogen by a fast soaking in a solution with increasing concentration of cryoprotectant to a final of 16-18% (v/v) glycerol. The diffraction data were collected at ESRF synchrotron beam line ID30A-3 (Grenoble, France) at 100 K. Data processing was performed by the autoPROC toolbox (Vonrhein *et al*, 2011), which exploits XDS program for data reduction (Kabsch, 2010), the POINTLESS for space-group determination (Evans, 2011), and the AIMLESS program for scaling and merging (Akey *et al*, 2016), and STARANISO for analysis of diffraction anisotropy (Vonrhein *et al*, 2018).

### Crystal structure determination and refinement

The structure was solved by molecular replacement using the software Autorickshaw (Panjikar *et al*, 2005). After several attempts, the best template was obtained using the AlphaFold3.0 model of the middle and C-terminal domains (middle domain: residues 247-416; C domain: residues 417-645) (Abramson *et al*, 2024). All attempts using the structures of HtpG_Mtb_ homologues or the entire AlphaFold3.0 model were unsuccessful. The structural refinement was performed using the program Phenix (Adams *et al*, 2010). The program Coot was employed for density inspection and manual modifications of protein coordinates. All protein model pictures were produced with PyMOL (Rosignoli & Paiardini, 2022). Atomic coordinates have been deposited in the PDB with 9THZ identification code.

### Molecular modelling

The three-dimensional structural model of HtpG_Mtb_ complex with ATP was generated using the latest AlphaFold3.0 modelling server (https://www.alphafoldserver.com), which produced five ranked models. Model reliability was assessed using the predicted local distance difference test (pLDDT), the interface predicted Template Modelling (ipTM) score and the predicted Template Modelling (pTM) score. The pLDDT score is a per-residue score that ranges from 0 to 100, with higher values corresponding to greater confidence in the accuracy of the structural model. The pTM score (range 0 to 1) estimates how well the overall structure model matches the true structure, while the ipTM score (range 0 to 1) specifically measures the accuracy of the interfaces between proteins in a complex.

### Surface Plasmon Resonance

Assessments of binding kinetics and affinity between Toll-like receptors (TLR4 Cloud-Clone Corp. Catalog #: RPA753Hu01; TLR4/MD2, R&D biosystem, Catalog #: 3146-TM) and the full length HtpG_Mtb_ were performed at 25 °C using Biacore T200 (Cytiva, Uppsala, Sweden). The TLRs were stably immobilised on a CM5 Sensor Chip at a flow rate of 10 µL/min using standard amine-coupling protocols. No protein was injected over the reference surface. HBS-P+ buffer (10 mM Hepes, 150 mM NaCl and 0.05% tween 20) was used as a running buffer. All samples and buffers were degassed prior to use. Protein solutions in HBS-P+ buffer at various concentrations were injected at 25 °C with a flow rate of 30 μL/min for 120 s (association phase), and then the buffer alone was injected for 300 s (dissociation phase). Surface regeneration was performed by injecting glycine buffer at a flow rate of 30 µL/min (10 mM, pH 1.5, 1 min). All mathematical manipulations and fitting operations were performed using BIAevaluation software (v2.02) provided with the Biacore T200 instrument (Cytiva) and assuming a 1:1 Langmuir binding model and hetereogeneous ligand model.

### ATPase assay

ATP hydrolysis assays were performed using the PiPer Phosphate Assay kit (Invitrogen). Purified protein at 5 μM was incubated with different ATP concentration in a range of 0 –1600 μM at 37°C for 90-120 min. Fluorescence was measured at 544 nm/590 nm (excitation/emission) on a EnVision XCite Multimode Plate Reader (Revvity) and corrected by the following equation: enzyme activity = full reaction (all components) -no enzyme control (ATP background) - no substrate control (enzyme only) + no enzyme / no substrate control (buffer background). Fluorescence was converted to free phosphate using a phosphate standard curve. ATP activity was normalised respect protein concentration and data were analysed using the program Prism. Enzyme activities are averages of at least three independent measurements.

## Data availability

Structural data generated in this study have been deposited in the Protein Data Bank (PDB) under the accession code 9THZ.

## Author contributions

RB and AR conceived the project. GB produced recombinant proteins and performed crystallisation. AR conducted biochemical experiments. GB and RB performed structure determination. MS, MCS and PC performed SPR analysis. RB wrote the manuscript and analysed data. GB, AR, MS, MCS, HJK, RB revised the manuscript.

## Acknowledgements

We would like to acknowledge the kind help of Dr Santosh Panjikar in solving the crystal structure using Autorickshaw. We also acknowledge the European Synchrotron Radiation Facility for provision of beam time on ID30A-3. Funding was provided by the project INF-ACT “One Health Basic and Translational Research Actions addressing Unmet Needs on Emerging Infectious Diseases PE00000007”, PNRR Mission 4, EU “NextGenerationEU”- D.D. MUR Prot.n. 0001554 of 11/10/2022 and by the project EVADERE (Enhanced VAccines DEsign against antimicrobial Resistance) Marie Skłodowska-Curie Actions (2026-2030). A.R. and M.S. were funded by the project PRIN 2022 PNRR, Funded by the European Union, “NextGenerationEU”, Mission 4 Component 2. CUP B53D23033100001, Prot. P2022JE8FN- “FIGHT_TB:FIndinG High-grade anTigens towards an innovative TB vaccine” - D.D. MUR Prot. n. 0001363 of 1 September 2023.

